# GDF15 and the beneficial actions of metformin in pre-diabetes

**DOI:** 10.1101/677831

**Authors:** Anthony P Coll, Michael Chen, Pranali Taskar, Debra Rimmington, Satish Patel, John Tadross, Irene Cimino, Ming Yang, Paul Welsh, Samuel Virtue, Deborah A. Goldspink, Emily Miedzybrodzka, Y. C. Loraine Tung, Sergio Rodriguez-Cuenca, Rute A. Tomaz, Heather P. Harding, Audrey Melvin, Giles S.H. Yeo, David Preiss, Antonio Vidal-Puig, Ludovic Vallier, David Ron, Fiona M. Gribble, Frank Reimann, Naveed Sattar, David B. Savage, Bernard B. Allan, Stephen O’Rahilly

## Abstract

Metformin, the world’s most prescribed anti-diabetic drug, is also effective in preventing Type 2 diabetes in people at high risk, by lowering body weight, fat mass and circulating insulin levels through mechanisms that are incompletely understood. Recent observational studies reporting the association of metformin use and circulating levels of GDF15 led us to hypothesize that GDF15, which signals through a specific receptor complex in the hindbrain to reduce body weight, might mediate these effects. We measured GDF15 in people without diabetes from a randomized placebo-controlled trial of metformin. Over 18 months, participants allocated metformin lost significant weight and levels of GDF15 were persistently elevated compared to placebo. The change in plasma GDF15 in this study correlated positively with weight loss. In wild-type mice, oral metformin increased circulating GDF15 with *GDF15* expression increasing predominantly in the distal intestine and the kidney. Metformin prevented weight gain in response to high fat diet in wild-type mice but not in mice lacking GDF15 or its receptor GFRAL. In obese, high fat-fed mice, the effects of metformin to reduce body weight were reversed by a GFRAL antagonist antibody. Metformin had effects on both energy intake and energy expenditure that required GDF15. The insulin sensitising effects of metformin determined by insulin tolerance were abolished in mice lacking GDF15. Metformin significantly reduced fasting glucose and insulin levels in wild type but not in mice lacking GDF15. In summary, metformin increases the circulating levels of GDF15, which appears to be necessary for many of its actions as a metabolic chemopreventive agent.

---

Metformin has been used as a treatment for Type 2 diabetes since the 1950s (PMID: 28776081). More recent studies have shown that it can also prevent or delay the onset of Type 2 diabetes in people at high risk ^1 2^. When compared to those receiving placebo, at-risk individuals treated with metformin manifest a modest but consistent reduction in body weight, lower circulating glucose and insulin levels and enhanced insulin sensitivity ^3^. Although many mechanisms of action for the insulin sensitizing actions of metformin have been proposed ^4^, few if any of these would explain the beneficial weight loss promoting effects of the drug which continue to be recognised ^5^. A recent observational epidemiological study^6^ noted a robust association of metformin use with circulating levels of GDF15, a peptide hormone produced by cells responding to a wide range of stressors^7^. GDF15 acts through a receptor complex solely expressed in the hindbrain, through which it suppress food intake ^8-11^. We hypothesized that metformin’s effects to lower body weight and perhaps some of its other salutary effects in pre-diabetes might involve the elevation of circulating levels of GDF15.

We first measured circulating GDF15 levels at 6, 12 and 18 months in all available participants from the CAMERA clinical trial ^12^, a randomized placebo-control trial of metformin in people without diabetes but with a history of cardiovascular disease. In this study, metformin treated participants were reported to lose ∼3.5% of starting body weight with no significant change in weight in the placebo arm^12^. Metformin treatment was associated with significantly (p < 0.001) increased levels of circulating GDF15 at all three time points relative to placebo (**Fig 1A).** Furthermore, the change in serum GDF15 from baseline in metformin recipients was significantly correlated (r=-0.26, p=0.024) with the amount of weight loss (**Fig 1B**).

**Figure 1.**
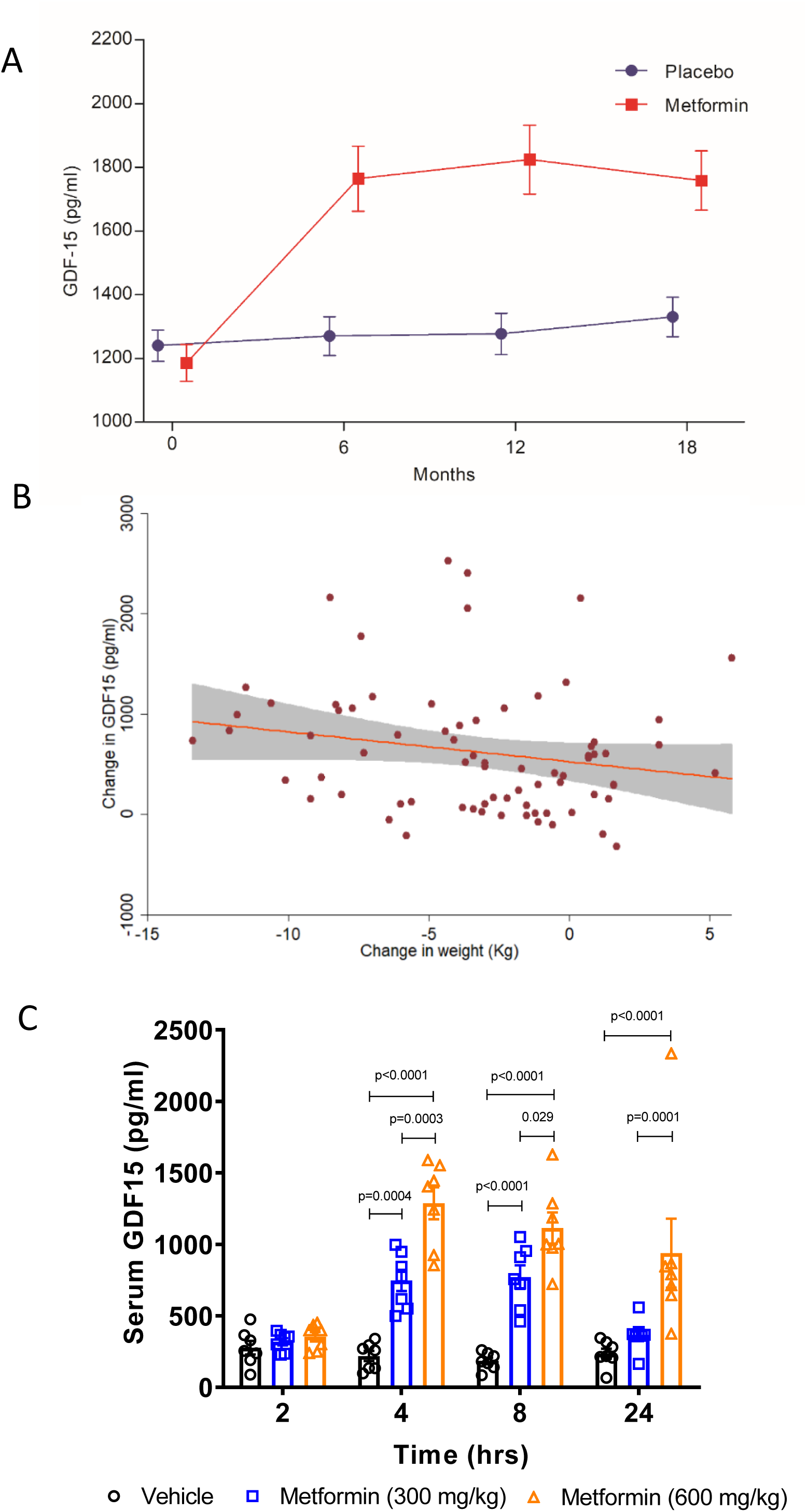
Effect of Metformin on circulating GDF15 levels in humans and mice. **A**) Plasma GDF15 concentration (mean± SEM) in overweight or obese non-diabetic participants with known cardiovascular disease randomised to metformin (n range =86) vs placebo (n range =85). **B**) Association between change in body weight and change in GDF15 among metformin treated participants (n=74, r=-0.26, p=0.024). Grey area is 95% confidence interval for slope. **C**)Serum GDF15 levels in obese mice measured 2, 4, 8 or 24 hours after a single oral dose of 300 mg/kg or 600 mg/kg metformin. Data are mean ± SEM (n=7/group); *P* by 2-way ANOVA with Tukey’s correction for multiple comparisons.

To determine if metformin treatment increased serum GDF15 levels in mice, we administered metformin to high fat fed mice by oral gavage and measured serum GDF15 levels. A single dose of 300 mg/kg of metformin increased GDF15 levels for at least 8 hours (**Fig 1C**). A higher dose of metformin, 600 mg/kg, increased serum GDF15 levels 4-6 fold at 4 and 8-hours post-dose, which were sustained over vehicle-treated mice for 24 hours. The effects of metformin on circulating GDF15 levels in lean, chow-fed mice were less pronounced (**Supplementary Fig 1**) suggesting an interaction between metformin and the high fat fed state.

In order to examine the extent to which the elevation in circulating GDF-15 induced by metformin is responsible for its effects on body weight, *Gdf15 +/+* and *Gdf15 -*^*/-*^ mice switched from chow to a high fat diet were dosed with metformin. High fat feeding induced similar weight gain in mice of both genotypes (**Fig 2A**). Metformin (300mg/kg/day), delivered by oral gavage starting 3 days after switching the diets, and maintained for 11 days, completely prevented diet-induced weight gain in *Gdf15* ^+/+^ mice but had no effect in *Gdf15* ^-/-^ mice (**Fig 2A**). In this study metformin significantly reduced cumulative food intake in wild type mice but this effect was abolished in *Gdf15*^-/-^ mice (**Fig 2B**). The identical protocol was applied to mice lacking GFRAL, the ligand-binding component of the hindbrain-expressed GDF15 receptor complex. Consistent with the results in mice lacking GDF15, metformin was unable to prevent weight gain in *Gfral* ^-/-^ mice (**Fig 2C**), despite similar levels of serum GDF15 **(Supplementary Fig 2A & B**). However, in this experiment, the reduction in cumulative food intake did not reach statistical significance (**Supplementary 2C**).

**Figure 2.**
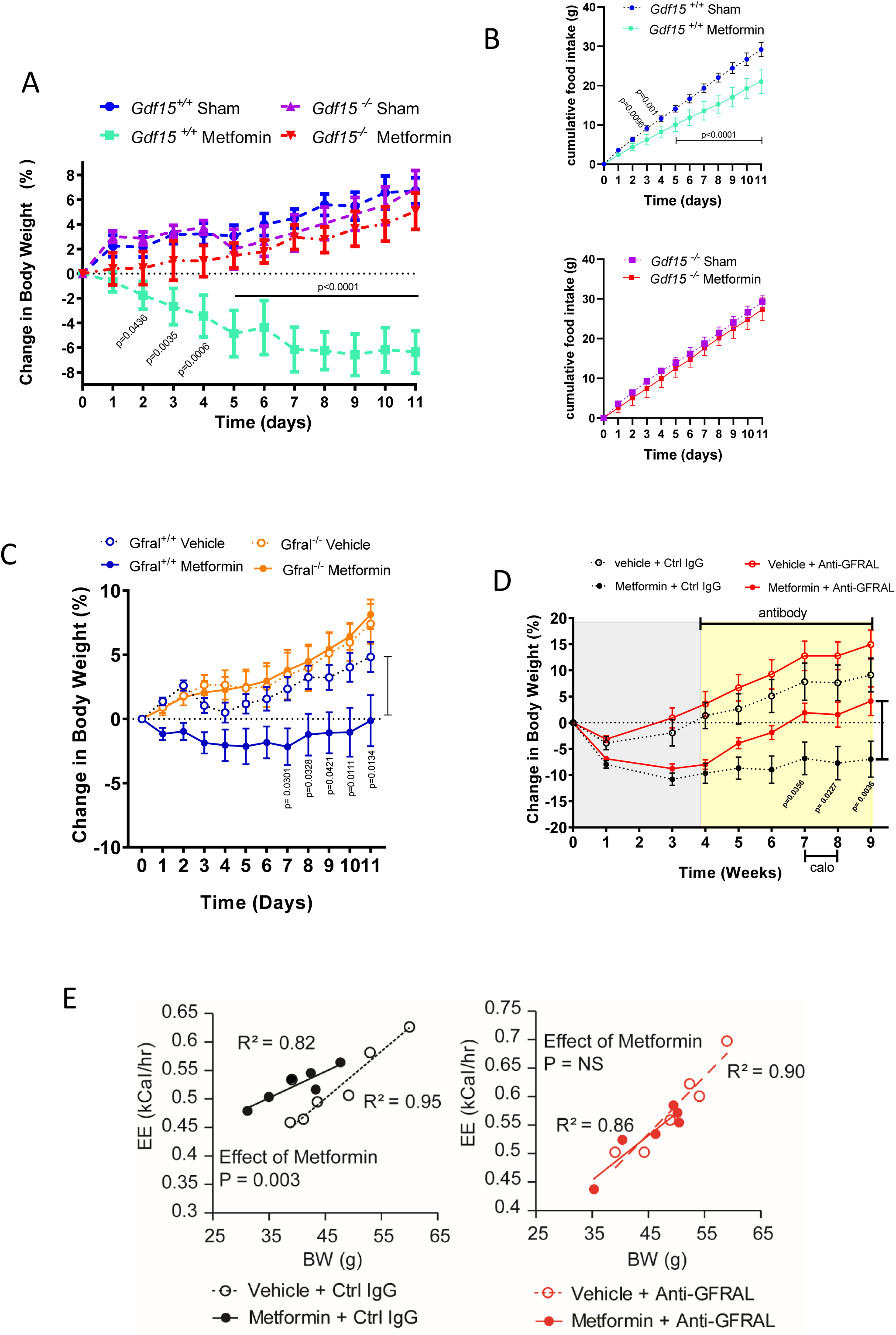
GDF15/GFRAL signalling is required for the weight loss effects of metformin on a high fat diet. **A**) Percentage change in body weight of *Gdf15+/+* and *Gdf15*^*-*^*/*^*-*^ mice on a high-fat diet treated with metformin (300mg/kg/day) for 11 days. Data are mean ± SEM (n=6/group except *Gdf15+/+* sham n=7); *P* by 2-way ANOVA with Tukey’s correction for multiple comparisons. **B) Cumulative** food intake of *Gdf15+/+* and *Gdf15*^*-*^*/*^*-*^ mice (upper and lower panel, respectively) on a high-fat diet treated with metformin (300mg/kg/day) for 11 days. Mice as in **Figure 2 A;** *P* by 2-way ANOVA with Sidak’s correction for multiple comparisons. **C)** Percentage change in body weight of *Gfral +/+* and *Gfral* ^*-*^*/*^*-*^ mice on a high-fat diet treated with metformin (300mg/kg/day) for 11 days. Data are mean ± SEM, n=6 all groups);*P* by 2-way ANOVA with Tukey’s correction for multiple comparisons. **D**) Percentage change in body weight of metformin-treated obese mice dosed with an anti-GFRAL antagonist antibody, weekly for 5 weeks (yellow), starting 4 weeks after initial metformin exposure (grey). Data are mean ± SEM (n=7 Veh + control IgG and Metformin + anti –GFRAL; n=8 other groups); *P* by 2-way ANOVA with Tukey’s correction for multiple comparisons. “calo” = period in which energy expenditure measured(**see Figure 2E**). **E**) Scatter plot of ANCOVA analysis of energy expenditure against body weight of mice treated with anti-GFRAL or control IgG (mice as in **Figure 2D**) 7 weeks after starting metformin, n=6 mice/group. Data are individual mice and *P* for metformin calculated using ANCOVA with body weight as a covariate and treatment as a fixed factor.

To investigate the contribution of GDF15/GFRAL signalling to sustained, metformin-dependent weight regulation, we performed a 9-week study in which mice received approximately 250-300 mg/kg/day of metformin that was incorporated into their high-fat diet. The mice lost approximately 10% body weight after 1 month on this diet (**Fig 2D**). At this time, we started dosing mice with an anti-GFRAL antagonist antibody, with a single dose administered every week. Metformin-consuming mice treated with anti-GFRAL gained ∼12% body weight after 5 weeks, while IgG control treated mice maintained a steady weight 10% below their starting weight (**Fig 2D**). The weight gain in anti-GFRAL treated mice was exclusively from increased fat mass (**Supplementary Fig 2D)**.

While GDF15 dependent effects on suppression of food intake clearly contributes to the weight loss seen with recombinant GDF15^8-11^ and with metformin, we undertook experiments to see if there were additional effects on energy expenditure by undertaking indirect calorimetry in metformin-and placebo-treated mice treated with anti-GFRAL antibody. All data were analysed by ANCOVA with body weight as the co-variate. Metformin treatment resulted in a statistically significant increase in metabolic rate which was blocked by antagonism of GFRAL (**Fig 2E**). We conclude that, under conditions where GDF15 levels are increased by metformin, body weight reduction is contributed to by both reduced food intake and an inappropriately high energy expenditure.

To examine the extent to which the insulin sensitising effects of metformin are dependent on GDF15 we repeated the experiment described in Fig 2A (see **Supplementary Fig 3**), this time undertaking insulin tolerance testing in metformin and sham-treated GDF15 null mice and their wild type littermates after 11 days of high fat feeding. In wild type mice, metformin significantly increased insulin sensitivity as assessed by the area under the plasma glucose curve at an ITT (**Fig 3A**). This effect was abolished in mice lacking GDF15.

**Figure 3.**
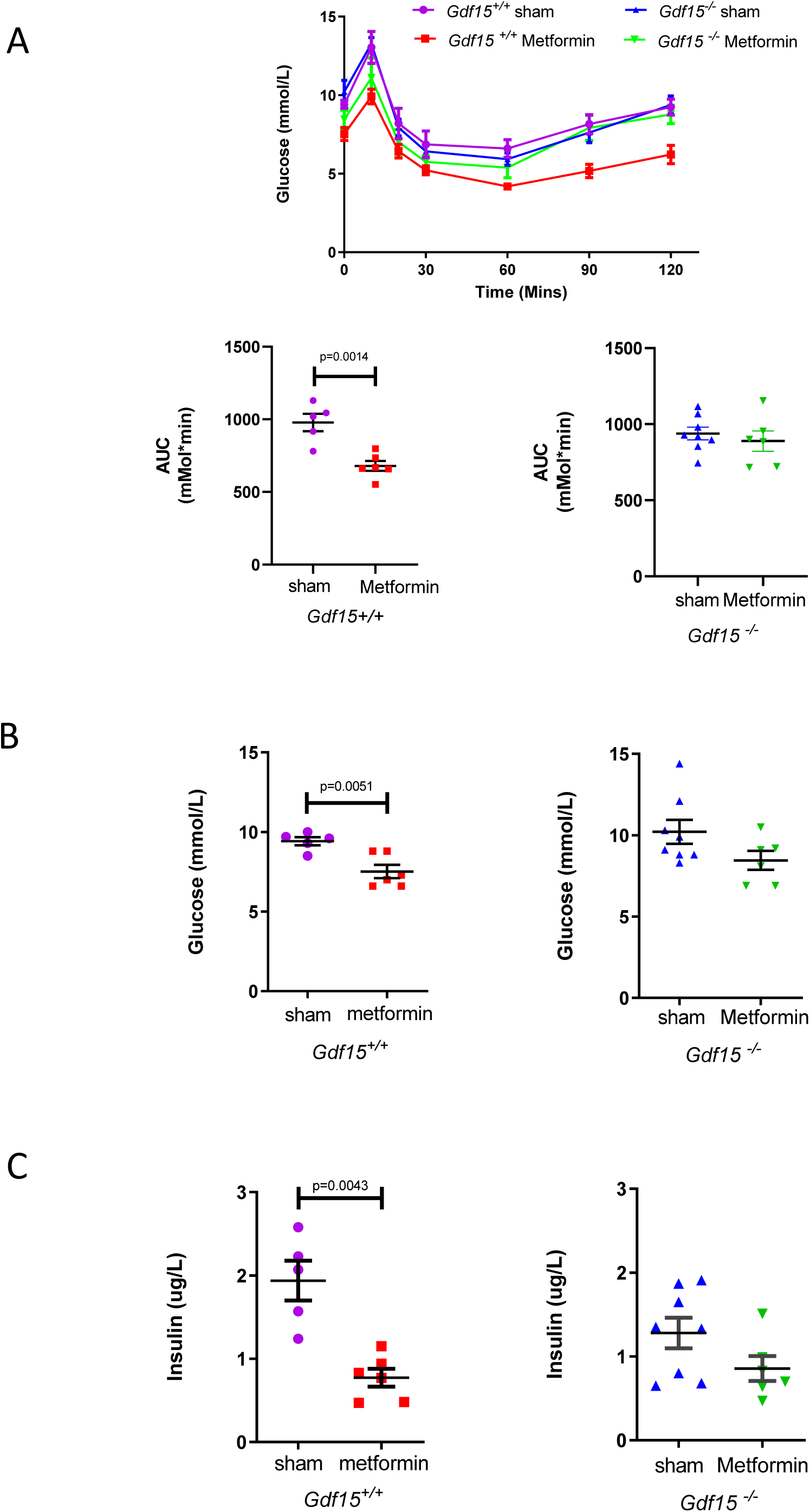
Effects of metformin on glucose homeostasis. **A**) Insulin tolerance test (intraperitoneal insulin (0.5 IU/kg) 4 hrs after last dose of metformin. High fat diet fed *Gdf15 +/+* and *Gdf15* ^*-*^*/*^*-*^ received oral dose of metformin (300mg/kg) once daily for 11 days. Data are mean ± SEM (n=6/group, except *Gdf15* ^-/-^ sham= 8, *Gdf15+/+* sham= 5). Upper panel; glucose levels over time after insulin administration: lower panels; area under the curve analysis, P by two tailed *t*-test. **B**) Plasma glucose and **C**) plasma Insulin at time 0 mins, mice as **Figure 3A**, *P* by two-tailed *t*-test (B) or Mann-Whitney test (C).

Both fasting blood glucose and fasting insulin were reduced by metformin with the effect of metformin being modulated by the presence or absence of GDF15. Metformin significantly (p<0.05) reduced mean fasting insulin by 60 % in wild type mice with a non-significant 33% reduction seen in mice lacking GDF15 (**Fig 3B, 3C**).

To establish which tissues were responsible for the rise in circulating GDF15 after metformin administration we examined gene expression in a tissue panel obtained from mice fed a high fat diet (for 4 weeks) and sacrificed 6 hours after a single gavage dose of metformin (600mg/kg). Circulating concentrations of GDF15 increased ∼4-fold compared to sham treated mice **(Supplementary Fig 4).** *Gdf15* mRNA was significantly increased by metformin in small intestine, colon and kidney after 6 hours, but not in skeletal muscle or adipose tissue **(Fig 4A).** In situ hybridisation studies demonstrated strong *Gdf15* expression in crypt enterocytes in the colon and small intestine and in periglomerular renal tubular cells. (**Fig 4B, Supplementary Fig 5**). We confirmed these sites of tissue expression in a separate cohort of HFD fed mice (those used in Fig 2A), treated with metformin for 11 days (**Supplementary Fig 6**).

**Figure 4.**
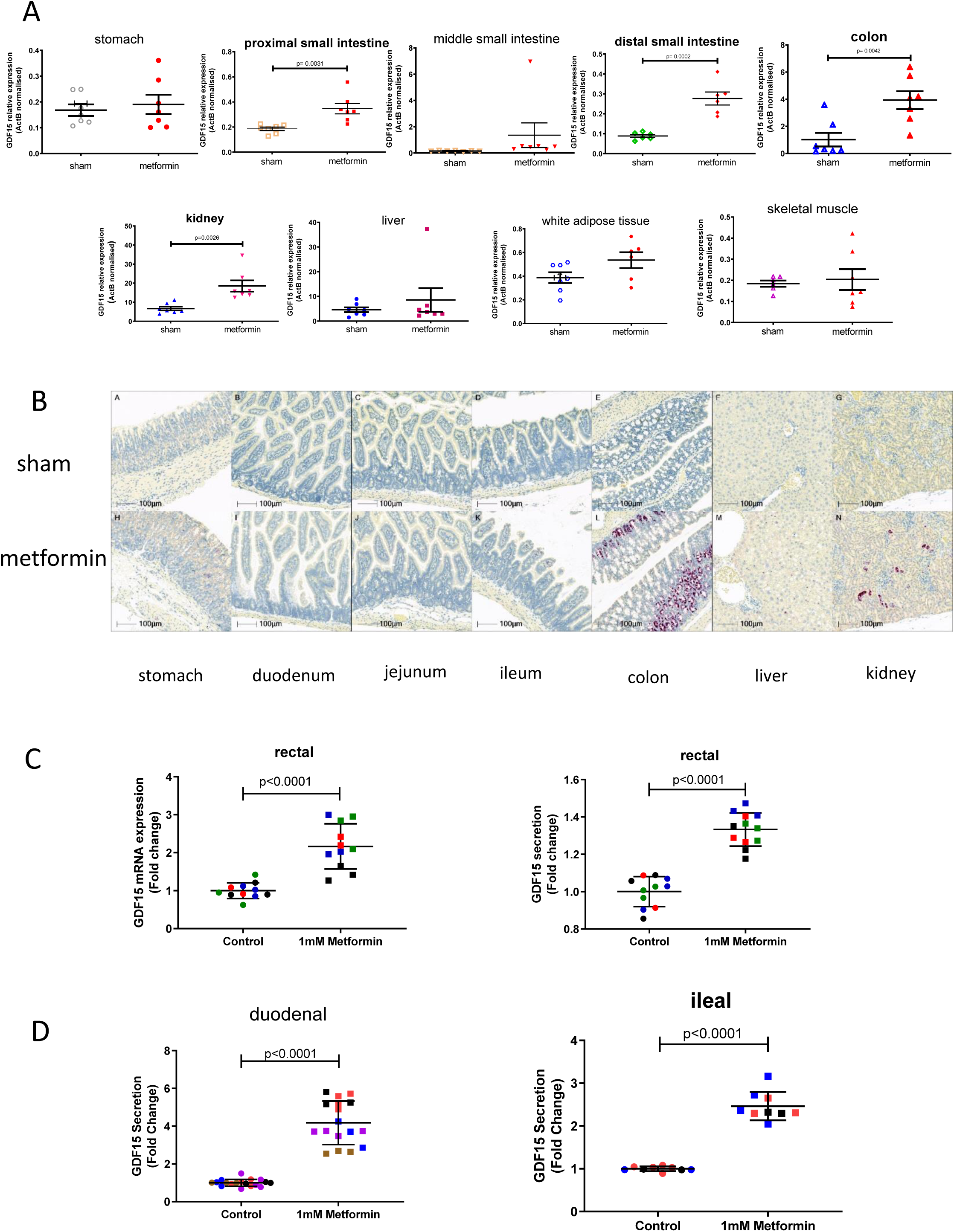
Metformin increases GDF15 expression in the enterocytes of distal intestine and the renal tubular epithelial cells. Analysis of *Gdf15* mRNA expression in tissues collected from high-fat fed wild type mice 6 hrs after single dose of oral metformin (600mg/kg). Data are mean ± SEM, n=7/group. **A**) mRNA analysis of fresh frozen tissue normalised to expression levels of ActB.*P* by two tailed *t*-test. **b**) In situ hybridization for *Gdf15* mRNA (red spots; n= 7 per group, representative images from the mouse with plasma GDF15 closest to group median). ***C****) Gdf15* mRNA expression (left) and GDF15 protein in supernatant (right) of human derived 2D monolayer rectal organoids treated with metformin. Each colour represents independent experiments (n= 4), data are mean ± SEM, *P* by two-tailed *t*-test. **D**) GDF15 protein in supernatants of mouse-derived 2D monolayer duodenal (left) and ileal (right) organoids treated with metformin. Each colour represents independent experiment (duodenal n= 5, ileal n=3), data are mean ± SEM, *P* by two-tailed *t*-test.

Further, in human (**Fig 4C**) and murine (**Fig 4D**) intestinal-derived organoids grown in 2D transwells and treated with metformin, we saw a significant induction of mRNA expression and GDF15 protein secretion.

Given the proposed importance of the liver for metformin metabolic action it was notable that GDF15 levels were only increased in the liver of a single mouse (Fig 4A). To test whether hepatocytes are capable of responding to biguanide drugs with an increase in GDF15 we incubated freshly isolated murine hepatocytes (**Supplementary Fig 7A**) and stem-cell derived human hepatocytes (**Supplementary Fig 7B**) with metformin and found a clear induction of GDF15 expression. Additionally, acute administration of the more cell penetrant biguanide drug phenformin to mice significantly increased circulating GDF15 levels (**Supplementary Fig 7C**) and markedly increased *Gdf15* mRNA expression in hepatocytes in a predominantly peri-central pattern (**Supplementary Fig 7D&E**).

GDF15 expression has been reported to be a downstream target of the cellular integrated stress response (ISR) pathway^13-15^.*Gdf15* mRNA levels were increased in kidney and colon 24 h after a single oral dose of metformin and these changes correlated positively with the fold elevation of CHOP mRNA **(Fig 5 A&B**). As phenformin has broader cell permeability than metformin^16^ we used it to explore the effects of biguanides on the ISR and its relationship to GDF15 expression in vitro. In murine embryonic fibroblasts (MEFs), which do not express the organic cation transporters needed for the uptake of metformin, phenformin (but not metformin) increased EIF2α phosphorylation, ATF4 and CHOP expression, **(Fig 5C)** and GDF15 mRNA (**Fig 5D**), though the changes in EIF2a phosphorylation and ATF4 and CHOP expression were modest compared with those induced by tunicamycin despite similar levels of GDF15 mRNA induction. Both genetic deletion of ATF4 and siRNA-mediated knockdown of CHOP in MEFs significantly reduced phenformin-mediated induction of GDF15 mRNA expression (**Fig 5E and 5F**). In addition, phenformin induction of GDF15 was markedly reduced by co-treatment with the EIF2α inhibitor, ISRIB but, notably, not by the PERK inhibitor, GSK2606414 (**Fig 5G**). Further, GDF15 secretion in response to metformin in murine duodenal organoids was also significantly reduced by co-treatment with ISRIB **(Fig 5H).** However, gut organoids derived from CHOP null mice are still able to increase GDF15 secretion in response to metformin (**Fig 5I**) indicating the existence of CHOP-independent pathways under some circumstances. The data suggest that the effects of biguanides on GDF15 expression are at least partly dependent on the ISR pathway but are independent of PERK. However, the relative importance of components of the ISR pathway may vary depending on specific cell type, dose and agent used.

**Figure 5.**
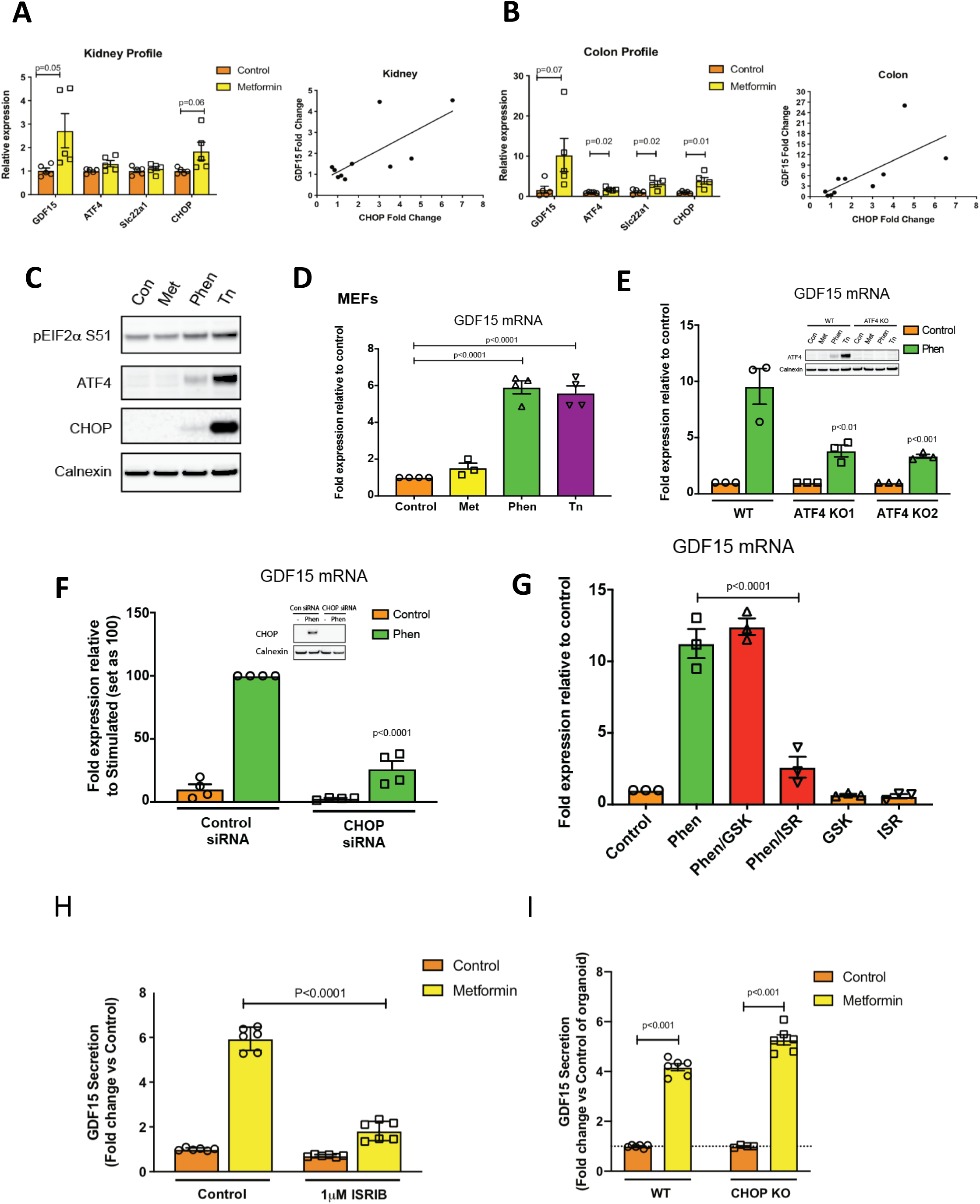
Role of the Integrated Stress Response (ISR) in biguanide-induced. ***Gdf15* expression** mRNA levels in kidney (**A**) and colon (**B**) isolated from obese mice 24 hours after a single oral dose of metformin (600mg/kg). Data are mean ± SEM (n=5/group). P values were calculated by two-tailed *t*-test. GDF15 mRNA fold induction 24 hrs post metformin 600mgs/kg is positively correlated with CHOP mRNA induction in both kidney (**A**, right panel) and colon (**B**, right panel). (**C**) Immunoblot analysis of ISR components and (**D**) GDF15 mRNA expression in wild type MEFs (mouse embryonic fibroblasts) treated with vehicle control (Con), metformin (Met, 2 mM) or phenformin (Phen, 5 mM) or tunicamycin (Tn, 5 μg/ml -used as a positive control) for 6 h. (**E**) GDF15 mRNA expression in ATF4 knockout (KO) MEFs or (**F**) in control siRNA and CHOP siRNA transfected WT MEFs treated with Tn or Phen for 6 h or (**G**) in WT MEFs pre-treated for 1 h either with the PERK inhibitor GSK2606414 (GSK, 200 nM) or eIF2α inhibitor ISRIB (ISR, 100 nM) then co-treated with Phen for a further 6 h. mRNA expression is presented as fold-expression relative to its respective control treatment (set at 1) or phen treated samples (set as 100) with normalisation to HPRT gene expression. Data is expressed as mean ± SEM from two for (D) and at least three independent experiments for (E-G). *P* by two tailed t-test. (**H**)GDF15 protein in supernatant of mouse derived 2D duodenal organoids treated with metformin in the absence or presence of ISRIB (1 μM). Data is expressed as mean ± SEM from two independent experiments, *P* by 2 way ANOVA with Sidak’s correction for multiple comparisons. (**I**)GDF15 protein in supernatants of mouse-derived 2D duodenal organoids from wild type and CHOP null mice treated with metformin from two independent experiments. Data are mean ± SEM, *P* by two-tailed t-test.

Our observations represent a significant advance in our understanding of pleiotropic actions of metformin, one of the world’s most frequently prescribed drugs. Metformin has recently been shown to increase circulating GLP1 levels^17-19^, but its metabolic effects in mice are unimpaired in mice lacking the GLP-1 receptor ^20^. Metformin has also been reported to alter the intestinal microbiome^21,22^ but it is challenging to design experiments to firmly establish the causal relationship of that change to the beneficial effects of the drug ^23^.

In the work presented herein, we describe a body of data from humans, cells, organoids and mice that securely establish a role for GDF15 in the mediation of metformin’s effects in the pre-diabetic state. Future studies will explore the extent to which the GDF15 dependent effects on circulating insulin and insulin sensitivity are mediated by its effects on energy balance or whether GDF15 has additional independent metabolic actions. The extent to which metformin’s glucose lowering actions in Type 2 diabetes are mediated by GDF15 will also require additional study.

It is notable that the lower small intestine and colon are a major site of metformin induced GDF15 expression. A body of work is emerging which strongly implicates the intestine as a major site of metformin action, with metformin being demonstrated to cause increased glucose uptake into colonic epithelium from the circulation^24^ and with a gut-restricted formulation of metformin showing enhanced glucose lowering efficacy over formulations which are systemically absorbed^25^. Our finding that the intestine is a major site of metformin-induced GDF15 expression provides a further mechanism through which metformin action on the intestinal epithelium may mediate a substantial proportion of its beneficial metabolic effects.

## Acknowledgments

CAMERA trial funded by a project grant from the Chief Scientist Office, Scotland (CZB/4/613) D.P. supported by a University of Oxford British Heart Foundation Centre of Research Excellence Senior Transition Fellowship (RE/13/1/30181).

N.S. and P.W. acknowledge support from BHF Centre of Excellence award (COE/RE/18/6/34217).

The authors would like to thank Peter Barker, Keith Burling and other members of the Cambridge Biochemical Assay Laboratory (CBAL). This project is supported by the National Institute for Health Research (NIHR) Cambridge Biomedical Research Centre. The views expressed are those of the authors and not necessarily those of the NIHR or the Department of Health and Social Care.

A.P.C., D.Rimmington, J.T., I.C., Y.C.L.T. and G.S.H.Y. are supported by the Medical Research Council (MRC Metabolic Diseases Unit [MC_UU_00014/1])

Mouse studies in Cambridge supported by Sarah Grocott and other members of the Disease Model Core, part of the MRC Metabolic Diseases Unit (MC_UU_00014/5) and Wellcome Trust Strategic Award (100574/Z/12/Z).

D.B.S. and S.O.R. are supported by the Wellcome Trust (WT 107064 and WT 095515/Z/11/Z), the MRC Metabolic Disease Unit (MC_UU_00014/1), and The National Institute for Health Research (NIHR) Cambridge Biomedical Research Centre and NIHR Rare Disease Translational Research Collaboration.

We thank Julia Jones and other members of Histopathology and ISH Core Facility, Cancer Research UK Cambridge Institute, University of Cambridge, Li Ka Shing Centre, Robinson Way, Cambridge CB2 0RE, UK.

D. Ron is supported by a Wellcome Trust Principal Research Fellowship (Wellcome 200848/Z/16/Z) and a Wellcome Trust Strategic Award to the Cambridge Institute for Medical Research (Wellcome 100140).

A.V.-P., S.R.-C.and S.V. are supported by the BHF (RG/18/7/33636) and MRC (MC_UU_00014/2).

A.M. is supported by a studentship from the Experimental Medicine Training Initiative/AstraZeneca.

R.A.T. and L.V. are supported by ERC advanced grant NewChol and core support from the Wellcome Trust and Medical Research Council to the Wellcome–Medical Research Council Cambridge Stem Cell Institute.

M.Y., D.A.G., E.M., F.M.G. and F.R. are supported by the MRC (MC_UU_00014/3) and Wellcome Trust (106262/Z/14/Z and 106263/Z/14/Z). M.Y. is supported by a BBSRC-DTP studentship. CHOP null mice were kind gift of Dr Jane Goodall (University of Cambridge).

## Author Contributions

Overall conceptualization of studies included in this body of work by A.P.C., N.S., D.B.S., B.B.A. and S.O’R.

A.P.C., M.C., P.T., D.Rimmington, I.C. and Y.C.L.T. designed, managed, performed and analysed data from mouse experiments.

S.V. designed experiments and analysed data.

A.M. and G.S.H.Y. contributed to conceptualisation of experiments and data analysis.

J.T. performed ISH experiments.

S.P. designed, managed and performed cell based assays along with E.M., S.R.C., R.A.T., H.P.H., A.V-P., L.V. and D.Ron.

M.Y., D.A.G., F.M.G., F.R. designed, performed and analysed organoid experiments.

P.W., D.P. and N.S. designed, analysed and interpreted data arising from the CAMERA study.

A.P.C., D.B.S., B.B.A. and S.O.R. wrote the paper, which was reviewed and edited by all the authors.

## Methods

### Human Studies

CAMERA was a randomized, double-blinded, placebo-controlled trial designed to investigate the effect of metformin on surrogate markers of cardiovascular disease in patients without diabetes, aged 35 to 75, with established coronary heart disease and a large waist circumference (≥ 94cm in men, ≥80 cm in women) (NCT00723307). This single-centre trial enrolled 173 adults who were followed up for 18 months each. A detailed description of the trial and its results has been published previously^12^. In brief, participants were randomized 1:1 to 850mg metformin or matched placebo twice daily with meals. Participants attended six monthly visits after overnight fasts and before taking their morning dose of metformin. Blood samples collected during the trial were centrifuged at 4 degrees Celsius soon after sampling, separated and stored at -80°C

All participants provided written informed consent. The study was approved by the Medicines and Healthcare Products Regulatory Agency and West Glasgow Research Ethics Committee, and done in accordance with the principles of the Declaration of Helsinki and good clinical practice guidelines.

Serum GDF15 assays were completed by the Cambridge Biochemical Assay Laboratory, University of Cambridge. Measurements were undertaken with antibodies & standards from R&D Systems (R&D Systems Europe, Abingdon UK) using a microtiter plate-based two-site electrochemiluminescence immunoassay using the MesoScale Discovery assay platform (MSD, Rockville, Maryland, USA).

### Mouse Studies

Studies were carried out in two sites; NGM Biopharmaceuticals, California, USA and Combined Facility, University of Cambridge, UK.

At NGM, all experiments were conducted with NGM IACUC approved protocols and all relevant ethical regulations were complied with throughout the course of the studies, including efforts to reduce the number of animals used. Experimental animals were kept under controlled light (12hour light and 12hour dark cycle, dark 6:30 pm -6:30 am), temperature (22 ± 3°C) and humidity (50% ± 20%) conditions. They were fed *ad libitum* on 2018 Teklad Global 18% Protein Rodent Diet containing 24 kcal% fat, 18 kcal% protein and 58 kcal% carbohydrate, or on high fat rodent diet containing 60 kcal% fat, 20 kcal% protein and 20 kcal% carbohydrates from Research Diets D12492i,(New Brunswick NJ 089901 USA) herein referred to as “60%HFD”.

In Cambridge, all mouse studies were performed in accordance with UK Home Office Legislation regulated under the Animals (Scientific Procedures) Act 1986 Amendment, Regulations 2012, following ethical review by the University of Cambridge Animal Welfare and Ethical Review Body (AWERB). They were maintained in a 12-hour light/12-hour dark cycle (lights on 0700–1900), temperature-controlled (22°C) facility, with *ad libitum* access to food (RM3(E) Expanded chow, Special Diets Services, UK) and water. Any mice bought from an outside supplier were acclimatised in a holding room for at least one week prior to study. During study periods they were fed ad libitum high fat diet, either D12451i (45 kcal% fat, 20 kcal% protein and 35 kcal% carbohydrates, herein referred to as “45%HFD”) or D12492i (Research Diets, as above) as highlighted in individual study.

Sample sizes were determined on the basis of homogeneity and consistency of characteristics in the selected models and were sufficient to detect statistically significant differences in body weight, food intake and serum parameters between groups. Experiments were performed with animals of a single gender in each study. Animals were randomized into the treatment groups based on body weight such that the mean body weights of each group were as close to each other as possible, but without using excess number of animals. No samples or animals were excluded from analyses. Researchers were not blinded to group allocations.

### Mouse study 1. Acute two-dose metformin study in high fat diet fed mice

Single housed male mice fed 60% HFD were used. Metformin (Sigma-Aldrich # 1396309) was reconstituted in water at 30 mg/ml for oral gavage and given in early part of light cycle. Terminal blood was collected by cardiac puncture into EDTA-coated tubes. GDF15 levels were measured using Mouse/Rat GDF15 Quantikine ELISA Kit (Cat#: MGD-150, R&D Systems, Minneapolis, MN) according to the manufacturers’ instructions. RNA was isolated from tissues using the Qiagen RNeasy Kit. RNA was quantified and 500ng was used for cDNA synthesis (SuperScript VILO 11754050 ThermoFisher) followed by qPCR. All Taqman probes were purchased from Applied Biosystems. All genes are expressed relative to 18s control probe and were run in triplicate.

### Mouse study 2. Acute metformin study in chow fed animals

#### 2.i) *ad libitum* group

Male C57BL6/J mice (Charles River, Margate, UK) were studied at 11 weeks old. 500mg of metformin was dissolved in 20 mls of water to make a working stock of 25mg/ml. 1 hr after onset of light cycle mice received a single dose by oral gavage of either metformin at 300mg/kg dose (Sigma, PHR1084-500MG) or matched volume of sham (water). Weight (mean± SEM) of control and treatment groups were 27.2 ± 0.3 *vs* 26.7 ± 0.2 g, respectively on day of study. After gavage mice were returned to an individual cage and were sacrificed at relevant time point by terminal anaesthesia (Euthatal by Intraperitoneal injection). Blood was collected into Sarstedt Serum Gel 1.1ml Micro Tube, left for 30mins at room temperature, spun for 5mins at 10k at 4^0^C before being frozen and stored at -80^°^C until assayed. Mouse GDF15 levels were measured using a Mouse GDF15 DuoSet ELISA (R&D Systems) which had been modified to run as an electrochemiluminescence assay on the Meso Scale Discovery assay platform.

#### 2.ii) fasted group

Mice, conditions and methods as in (2.i) except male mice studied at 9 weeks old and that 12 hr prior to administration of metformin mice and bedding were transferred to new cages with no food in hopper. Weight (mean± SEM) of groups at gavage were 22.3±0.5 g and 23.2±0.7g for control and treatment groups, respectively.

### Mouse study 3. Metformin to high fat diet fed *Gdf15* ^*-/-*^ mice and wild type controls

C57BL/6N-Gdf15tm1a(KOMP)Wtsi/H mice (herein referred to as “*Gdf15* ^*-/-*^ mice”) were obtained from the MRC Harwell Institute which distributes these mice on behalf of the European Mouse Mutant Archive (www.infrafrontier.eu). The MRC Harwell Institute is also a member of the International Mouse Phenotyping Consortium (IMPC) and has received funding from the MRC for generating and/or phenotyping the C57BL/6N-Gdf15tm1a(KOMP)Wtsi/H mice. The research reported in this publication is solely the responsibility of the authors and does not necessarily represent the official views of the Medical Research Council. Associated primary phenotypic information may be found at www.mousephenotype.org. Details of the alleles have been published^26-28^.

Experimental cohorts of male *Gdf15* ^*-/-*^ and wild type mice were generated by het x het breeding pairs. Mice were aged between 4.5 and 6.5 months. Body weight of study groups (mean±SEM) were 38.2±1.0g; 38.8±0.6g for wild type sham and metformin treatment respectively, and 37.9±0.8g; 37.0±1.4g for Gdf15^-/-^ sham and metformin treatment respectively. One week prior to study start mice were single housed and 3 days prior to first dose of metformin treatment, mice were transferred from standard chow to 60% high fat diet. Each mouse received a daily gavage of either sham or metformin for 11 days, and their body weight and food intake measured daily in the early part of the light cycle. One data point of 25 food intake points collected on day11 of study was lost due to technical error (mouse; *Gdf15* ^+/+^ metformin). On day 11 mice were sacrificed by terminal anaesthesia 4 hours post gavage, blood was obtained as in study 2. Tissues were fresh frozen on dry ice and kept at -80^0^C until day of RNA extraction.

### Mouse study 4. Metformin to high fat diet fed *Gfral* ^*-/-*^ mice

*Gfral*^*-/-*^ mice were purchased from Taconic (#TF3754) on a mixed 129/SvEv-C57BL/6 background and backcrossed for 10 generations to >99% C57BL/6 background at NGM’s animal facility. Experimental cohorts were generated by het X het breeding pairs. Study design as Study 3, except terminal blood was collected into EDTA-coated tubes.

### Mouse study 5. Anti GFRAL antibody to metformin treated high fat diet fed mice

#### Anti-GFRAL antibody generation

Anti-GFRAL monoclonal antibodies were generated by immunizing C57Bl/6 mice with recombinant purified GFRAL ECD-hFc fusion protein, which was purified via sequential protein-A affinity and size exclusion chromatography (SEC) techniques using MabSelect SuRe and Superdex 200 purification media respectively (GE Healthcare), as described elsewhere (Suriben et al. under review 2019). An in-house pTT5 hIgK hIgG1 expression vector was engineered to include the DEVDG (caspase-3) proteolytic site N-terminal to the Fc domain. The heavy chains of anti-GFRAL mAbs were subcloned via EcoR1/HindIII sites of in-house engineered pTT5 hIgK hIgG1 caspase-cleavable vector.

Light chains of anti-GFRAL mAbs were also subcloned within the EcoR1/HindIII sites in the pTT5 hIgK hKappa vector. The antibody were transiently expressed in Expi293 cells (Thermo Fisher Scientific) transfected with the pTT5 expression vector, and purified from conditioned media by sequential protein-A affinity and size-exclusion chromatographic (SEC) methods using MabSelect SuRe and Superdex 200 purification media respectively (GE Healthcare). All purified antibody material was verified endotoxin-free and formulated in PBS for in vitro and in vivo studies. Characterization of anti-GFRAL functional blocking antibodies was carried out using a cell-based RET/GFRAL luciferase gene reporter assays, in vitro binding studies (ELISA and Biacore) and in vivo studies, as described elsewhere (Suriben et al. under review 2019). In all studies with anti-GFRAL, purified recombinant non-targeting IgG on the same antibody framework was used as control. Metformin was mixed with food paste made from the 60 kcal% fat diet (Research diet# D12492) using a food blender at a concentration to achieve an approximate consumption of 300mg/kg metformin per day per mouse. Male animals were single housed throughout and at start of study period body weight (mean ±SEM) was 43.7±1.4g, 42.3±1.4g, 41.9±1.1g,43.3±1.3g, veh + control IgG, veh +anti-GFRAL, metformin + control IgG, Metformin + anti-GFRAL, respectively.

Recombinant antibodies were administered by subcutaneous injection in the early part of the light cycle. Body composition (lean and fat mass) was analyzed by ECHO MRI M113 mouse system (Echo Medical Systems). The metabolic parameters oxygen consumption (VO2) and carbon dioxide production (VCO2) were measured by an indirect calorimetry system (LabMaster TSE System, Germany) in open circuit sealed chambers. Measurements were performed for the dark (from 6pm to 6am) or light (from 6am to 6pm) period under *ad libitum* feeding conditions. Mice were placed in individual metabolic cages and allowed to acclimate for a period of 24 hours prior to data collection in every 30 minutes.

### Mouse study 6. Insulin tolerance test after metformin treatment to high fat diet fed

***Gdf15***^***-/-***^ **and wild type controls.**

Mice generation and protocol as Study 3, except aged 4 to 6 months. Body weight (mean±SEM) of study groups were 32.8+/-1.0g; 33.1+/-1.4g for wild type Sham and Metformin treatment respectively, and 32.9+/-0.9g; 32.2+/-1.3g for *Gdf15*^*-/-*^ Sham and Metformin treatment respectively. On day 11, after final dose of metformin mice were fasted for 4 hours. Baseline venous blood sample was collected into heparinised capillary tube for insulin measurement and blood glucose was measured using approximately 2 μl blood drops using a glucometer (AlphaTrak2; Abbot Laboratories) and glucose strips (AlphaTrak2 test 2 strips, Abbot Laboratories, Zoetis). Mice were given intraperitoneal injection of insulin (0.5U/kg mouse, Actrapid, NovoNordisk Ltd) and serial mouse glucose levels measured at time points indicated. Mice were sacrificed by terminal anaesthesia as in Study 2. Mouse insulin was measured using a 2-plex Mouse Metabolic immunoassay kit from Meso Scale Discovery Kit (Rockville, MD, USA), performed according to the manufacturer’s instructions and using calibrators provided by MSD.

### Mouse study 7. Acute single high dose metformin study in high fat diet fed wild type mice

Male C57BL6/J mice (Charles River, Margate, UK) aged 14 weeks were switched from standard chow to 45 %HFD fat for 1 week then 60%HFD for 3 weeks (D12451i and D12492i, respectively, as above). At time of study (18 weeks old) body weights (mean ±SEM) were 40.4± 1.2g *vs* 41.1±1.3g, sham *vs* metformin group, respectively. 500mg of metformin (Sigma, PHR1084-500MG) was dissolved in 8.35 mls of water to make a working stock of 60mg/ml. Mice received a single dose by oral gavage of either 600mg/kg metformin or matched volume of sham (water). They were returned to ad lib 60 % fat diet and 6 hrs later blood was collected as study 2. Tissue samples for RNA analysis were collected into Lysing Matrix D homogenisation tube (MP Biomedicals) on dry ice and stored at -80^0^C until processed. Regions of gut were located and defined as per figure below;intestine between pylorus of stomach and caecum was laid out into 3 equal parts, with tissue taken from mid-point of each third (adapted from ^29^). Colon section was from mid-point between caecum and anus.

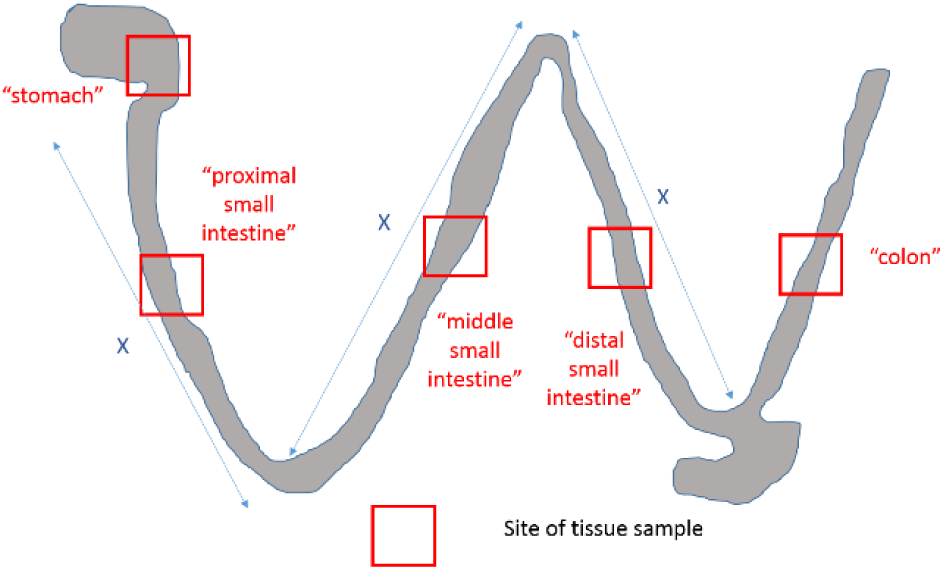

Tissue for in-situ hybridisation were dissected and placed into 10% formalin/PBS for 24hr at room temp, transfe
rred to 70% ethanol, and processed into paraffin. 5μm sections were cut and mounted onto Superfrost Plus (Thermo-Fisher Scientific). Detection of Mouse Gdf15 was performed on FFPE sections using Advanced Cell Diagnostics (ACD) RNAscope^®^ 2.5 LS Reagent Kit-RED (Cat No. 322150) and RNAscope^®^ LS 2.5 Probe Mm-Gdf15-O1 (Cat No. 442948) (ACD, Hayward, CA, USA). Briefly, sections were baked for 1 hour at 60oC before loading onto a Bond RX instrument (Leica Biosystems). Slides were deparaffinized and rehydrated on board before pre-treatments using Epitope Retrieval Solution 2 (Cat No. AR9640, Leica Biosystems) at 95°C for 15 minutes, and ACD Enzyme from the LS Reagent kit at 40oC for 15 minutes. Probe hybridisation and signal amplification was performed according to manufacturer’s instructions. Fast red detection of mouse Gdf15 was performed on the Bond RX using the Bond Polymer Refine Red Detection Kit (Leica Biosystems, Cat No. DS9390) according to the ACD protocol. Slides were then counterstained with haematoxylin, removed from the Bond RX and were heated at 60oC for 1 hour, dipped in Xylene and mounted using EcoMount Mounting Medium (Biocare Medical, CA, USA. Cat No. EM897L).

Slides imaged on an automated slide scanning microscope (Axioscan Z1 and Hamamatsu orca flash 4.0 V3 camera) using a 20x objective with a numerical aperture of 0.8.

Hybridisation specificity was confirmed by the absence of staining in Gdf15-/-mice.

RNA extraction was carried out with approximately 100mg of tissue in 1ml Qiazol Lysis Reagent (Qiagen 79306l) using Lysing Matrix D homogenisation tube and Fastprep 24 Homogeniser (MP Biomedicals) and Qiagen RNeasy Mini kit (Cat no 74106) using manufacturers’ protocols. 500ng of RNA was used to generate cDNA using Promega M-MLV reverse transcriptase followed by TaqMan qPCR in triplicates for GDF15. Samples were normalised to Act B. TaqMan Probes: Mm00442228 m1 GDF15, Mm02619580_g1 Act B, TaqMan;2X universal PCR Master mix (Applied Biosystems Thermo Fisher 4318157); QuantStudio 7 Flex Real time PCR system (Applied Biosystems Life Technologies)

### Mouse study 8. Acute phenformin study in standard chow-fed wild type animals

Male C57BL6/J mice aged 14 weeks with supplier, protocol and methods as study 2, except phenformin (Sigma PHR1573-500mg) used instead of metformin.

#### Organoid studies

Duodenal and ileal mouse organoid line generation, maintenance and 2D culture was performed as previously described^30^. CHOP null mice were kind gift of Dr Jane Goodall (University of Cambridge), with line from Jackson Laboratory, Maine (B6.129S(Cg)-Ddit3tm2.1Dron/J, Stock No: 005530) Human rectal organoids (experiments approved by the Research Ethics Committee under license number 09/H0308/24) were generated from fresh surgical specimens (Tissue Bank Addenbrooke’s Hospital (Cambridge, UK)) following a modified protocol ^30,31^. Briefly rectal tissue was chopped into 5mm fragments and incubated in 30 mM EDTA for 3x10mins, with tissue shaken in PBS after each EDTA treatment to release intestinal crypts. The isolated crypts were then further digested using TrypLE (Life Technologies) for 5 mins at 37°C to generate small cell clusters. These were then seeded into basement membrane extract (BME, R&D technology), with 20 μl domes polymerised in multiwell (48) dishes for 30-60 mins at 37°C. Organoid medium (Sato et al 2011) was then overlaid and changed 3 times per week. Human organoids were passaged every 14-21 days using TrypLE digestion for 15 mins at 37°C, followed by mechanical shearing with rigorous pipetting to breakup organoids into small clusters which were then seeded as before in BME. For transwell experiments TrypLE digested organoids were seeded onto matrigel (Corning) coated (2% for 60 mins at 37°C) polyethylene Terephthalate cell culture inserts, pore size 0.3 μm (Falcon) in organoid medium supplemented with Y-27632 (R&D technology). Organoids were observed through the transparent cell inserts to ensure 2D culture formation (allowing apical cell access for drug treatments). Medium was changed after 2 days and then switched on day 3 to a differentiation medium with wnt3A conditioned medium reduced to 10% and SB202190 / nicotinamide omitted from culture for 5 days.

For GDF 15 secretion experiments 2D cultured organoid cells were treated for 24 hrs with indicated drugs, with medium then collected and GDF15 measured at the Core Biochemical Assay Laboratory (Cambridge) using the human or mouse GDF15 assay kit as outlined in CAMERA human study and mouse study 2 above.

RNA was extracted using TRI reagent (Sigma), with any contaminated DNA eliminated using DNA free removal kit (Invitrogen). Purified RNA was then reverse transcribed using superscript II (Invitrogen) as per manufacturer’s protocol. RT-qPCR was performed on a QuantStudio 7 (Applied Biosystems) using Fast Taqman mastermix and the following probes (Applied Biosystems); Human GDF15 (Hs00171132_m1), Human ACTB (Hs01060665_g1). Gene expression was measured relative to β-actin in the same sample using the ΔCt method, with fold (cf. control) shown for each experiment.

#### Hepatocyte studies

### Primary mouse hepatocyte isolation and culture

Hepatocytes from 8-12 week old C57B6J male mice were isolated by retrograde, non-recirculating in situ collagenase liver perfusion. In brief: livers were perfused with modified Hanks medium without calcium (NaCl-8.0 g/L; KCl-0.4 g/L; MgSO4.7H2O-0.2 g/L; Na2HPO4.2H2O-0.12 g/L; KH2PO4-0.12 g/L; Hepes-3 g/L; EGTA-0.342 g/L; BSA-0.05 g/L) followed by digestion with perfusion media supplemented with calcium (CaCl2.2H2O-0.585 g/L) and 0.5mg/ml of collagenase IV (Sigma, C5138). The digested liver was removed and washed using chilled DMEM:F12 (Sigma) medium containing 2 mM L-glutamine, 10 % FBS, 1% penicillin/streptomycin (Invitrogen). Viable cells were harvested by Percoll (Sigma) gradient. The final pellet was resuspended in the same DMEM:F12 media. Cell viability was greater than 90%. Hepatocytes were plated onto primaria plates (Corning). Hepatocytes were allowed to recover and attach for 4-6 hr before replacement of the medium overnight prior to stress treatments the following day for the times and concentrations indicated.

### Generation and culture of iPSC derived human hepatocytes

The human induced pluripotent cell (hiPSC) line A1ATD^R/R^ used in this work was derived as previously described^32,33^, under approval by the regional research ethics committee (reference number 08/H0311/201). hiPSCs were maintained in Essential 8 chemically defined media^34 3^ supplemented with 2ng/ml Tgf-ß (R&D) and 25ng/ml FGF2 (R&D), and cultured on plates coated with 10µg/ml Vitronectin XF™ (STEMCELL Technologies). Colonies were regularly passaged by short-term incubation with 0.5mM EDTA in PBS. For hepatocyte differentiation, colonies were dissociated into single cells following incubation with StemPro™ Accutase™ Cell Dissociation Reagent (Gibco) for 5 minutes at 37°C. Single cell suspensions were seeded on plates coated with 10µg/ml Vitronectin XF™ (STEMCELL Technologies) in maintenance media supplemented with 10µM ROCK Inhibitor Y-27632 (Selleckchem) and grown for up to 72h prior to differentiation. Hepatocytes were differentiated as previously reported^35^, with minor modifications as listed. Briefly, following endoderm differentiation, anterior foregut specification was achieved after 5 days of culture with RPMI-B27 differentiation media supplemented with 50ng/ml Activin A (R&D)^35^. Foregut cells were further differentiated into hepatocytes with HepatoZYME-SFM (Gibco) supplemented with 2mM L-glutamine (Gibco), 1% penicillin-streptomycin (Gibco), 2% non-essential amino acids (Gibco), 2% chemically defined lipids (Gibco), 14μg/ml of insulin (Roche), 30μg/ml of transferrin (Roche), 50 ng/ml hepatocyte growth factor (R&D), and 20 ng/ml oncostatin M (R&D), for up to 27 days.

#### Cellular studies on integrated stress response

### Chemicals and Reagents

Tunicamycin and ISRIB were purchased from Sigma-Aldrich. Metformin and Phenformin was purchased from Cayman Chemicals and GSK2606414 from Calbiochem. The antibody for GDF15 and CHOP (sc-7351) were obtained from Santa Cruz. Phospho S51 EIF2 α (ab32157) and Calnexin (ab75801) were purchased from Abcam. The antibody for ATF4 was a kind gift from Dr David Ron (CIMR, Cambridge).

### Eukaryotic cell lines and treatments

Mouse embryonic fibroblast (MEF) cells lines were obtained from David Ron (CIMR/IMS, Cambridge) and maintained as previously described^15^. MEFs were transfected with 30 nM control siRNA or a smartpool on-target plus siRNA for mouse CHOP (Dharmacon - L-062068- 00-0005) using Lipofectamine RNAi MAX (Invitrogen) according to the manufacturer’s instruction. 48 h post siRNA transfection, cells were processed for RNA and protein expression analysis. All cells were maintained at 37 °C in a humidified atmosphere of 5 % CO2 and seeded onto 6- or 12-well plates prior to stress treatments for the times and concentrations indicated. Vehicle treatments (e.g. DMSO) were used for control cells when appropriate.

### RNA isolation/cDNA synthesis/Q-PCR

Following treatments, cells were lysed with Buffer RLT (Qiagen) containing 1 % 2-Mercaptoethanol and processed through a Qiashredder with total RNA extracted using the RNeasy isolation kit according to manufacturer’s instructions (Qiagen). RNA concentration and quality was determined by Nanodrop. 400 ng -500 ng of total RNA was treated with DNAase1 (Thermofisher Scientific) and then converted to cDNA using MMLV Reverse Transcriptase with random primers (Promega). Quantitative RT-PCR was carried out with either TaqMan™ Universal PCR Master Mix or SYBR Green PCR master mix on the QuantStudio 7 Flex Real time PCR system (Applied Biosystems). All reactions were carried out in either duplicate or triplicate and Ct values were obtained. Relative differences in the gene expression were normalized to expression levels of housekeeping genes, HPRT or GAPDH for cell analysis, using the standard curve method. Primers used for this study: mouse GDF15 (Mm00442228_m1 – ThermoFisher Scientific), human GDF15 (Hs00171132_m1 - ThermoFisher Scientific), human GAPDH (Hs02758991_g1 – ThermoFisher Scientific), mouse HPRT (Forward – AGCCTAAGATGAGCGCAAGT, reverse - GGCCACAGGACTAGAACACC)

### Immunoblotting

Following treatments, cells were washed twice with ice cold D-PBS and proteins harvested using RIPA buffer supplemented with cOmplete protease and PhosStop inhibitors (Sigma). The lysates were cleared by centrifugation at 13 000 rpm for 15 min at 4 °C, and protein concentration determined by a Bio-Rad DC protein assay. Typically, 20-30 μ g of protein lysates were denatured in NuPAGE 4× LDS sample buffer and resolved on NuPage 4-12 % Bis-Tris gels (Invitrogen) and the proteins transferred by iBlot (Invitrogen) onto nitrocellulose membranes. The membranes were blocked with 5 % nonfat dry milk or 5 % BSA (Sigma) for 1 h at room temperature and incubated with the antibodies described in the reagents section. Following a 16 h incubation at 4 °C, all membranes were washed five times in Tris-buffered saline-0.1% Tween-20 prior to incubation with horseradish peroxidase (HRP)-conjugated anti-rabbit immunoglobulin G (IgG), HRP-conjugated anti-mouse IgG (Cell Signalling Technologies). The bands were visualized using Immobilon Western Chemiluminescent HRP Substrate (Millipore). All images were acquired on the ImageQuant LAS 4000 (GE Healthcare).

### Statistical analyses

CAMERA data analysed using STATA version 15.1. Other statistical analyses were performed using Prism 7 and Prism 8, using unpaired 2 tailed t-tests or Mann-Whitney, or 2-way ANOVA, with multiple comparison adjustment by Tukey’s or Sidak’s test. Metabolic rate was determined using ANCOVA with energy expenditure as the dependent variable, body weight as a covariate and treatment as a fixed factor. ANCOVA was performed using SPSS 25 (IBM).

---

**Supplementary Figure 1.**
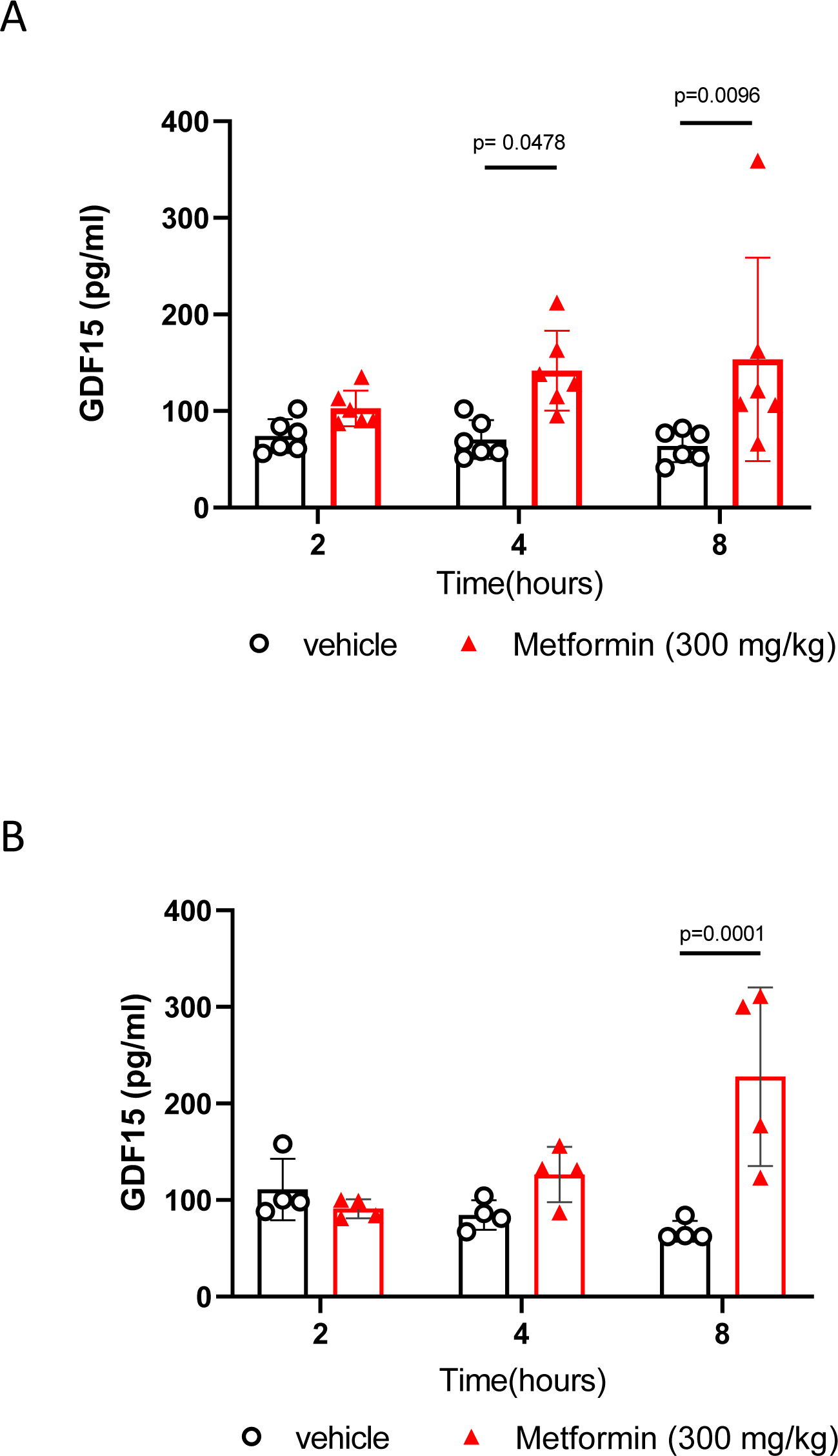
Effect of single oral dose of metformin in chow fed mice. Serum GDF15 levels in male mice measured 2, 4, or 8 hours after a single gavage dose of metformin (300mg/kg). **A**) mice *ad libitum* overnight fed prior to gavage. **B**) mice fasted for 12 hour prior to gavage. Data are mean ± SEM (A; n=6/group, B; n= 4/group); *P* by 2way ANOVA with Sidak’s correction for multiple comparisons.

**Supplementary Figure 2.**
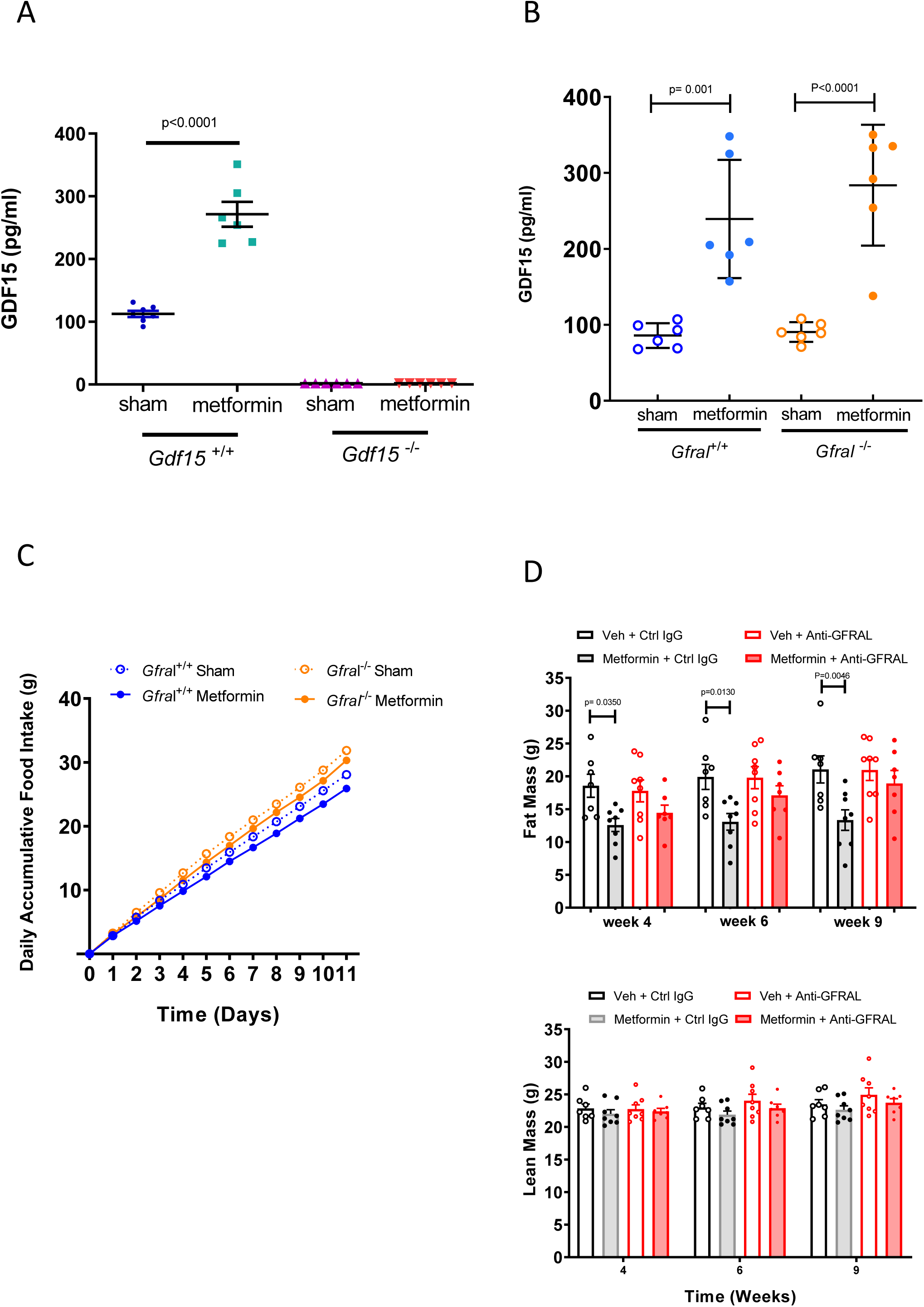
Response of high fat diet fed *Gdf15* ^*-/-*^ and *Gfral*^*-/*-^mice to metformin. **A**) Circulating GDF15 levels in high fat diet fed *Gdf15 +/+* and *Gdf15* ^*-*^*/*^*-*^ mice given oral dose of metformin (300mg/kg) once daily for 11 days. Samples were collected 4 hours after the final oral dose of metformin. Data are mean ± SEM, mice as in **Figure 2 A**. All samples from *Gdf15*^*-*^*/*^*-*^ were below lower limit of assay (< 2pg/ml), *P* by two-tailed *t*-test. **B**) Circulating GDF15 levels in high fat diet fed *Gfral +/+* and *Gfral* ^*-*^*/*^*-*^ mice given oral dose of metformin (300mg/kg) once daily for 11 days. Samples were collected 4 hours after the final oral dose of metformin. Data are mean ± SEM, mice as in **Figure 2C**, *P* by 2-way ANOVA with Tukey’s correction for multiple comparisons. **C**) Cumulative food intake in high fat diet fed *Gfral +/+* and *Gfral* ^*-*^*/*^*-*^ mice on a high fat diet given oral dose of metformin (300mg/kg) once daily for 11 days. Data are mean ± SEM, mice as in **Figure 2C**. **D**) Fat and lean mass in metformin-treated obese mice dosed with an anti-GFRAL antagonist antibody, weekly for 5 weeks, starting 4 weeks after initial metformin exposure (mice as **Figure 2D**). Body composition was measured using MRI after 4 weeks of metformin exposure, prior to receiving anti-GFRAL (week 4), after 6 weeks of metformin exposure and 2 weeks after receiving anti-GFRAL (week 6) and after 9 weeks of metformin exposure and 5 weeks after receiving anti-GFRAL (week 9). Data are mean ± SEM (n=7 Veh + control IgG and Metformin + anti – GFRAL; n=8 other groups); *P* by 2-way ANOVA with Sidak’s correction for multiple comparisons.

**Supplementary Figure 3.**
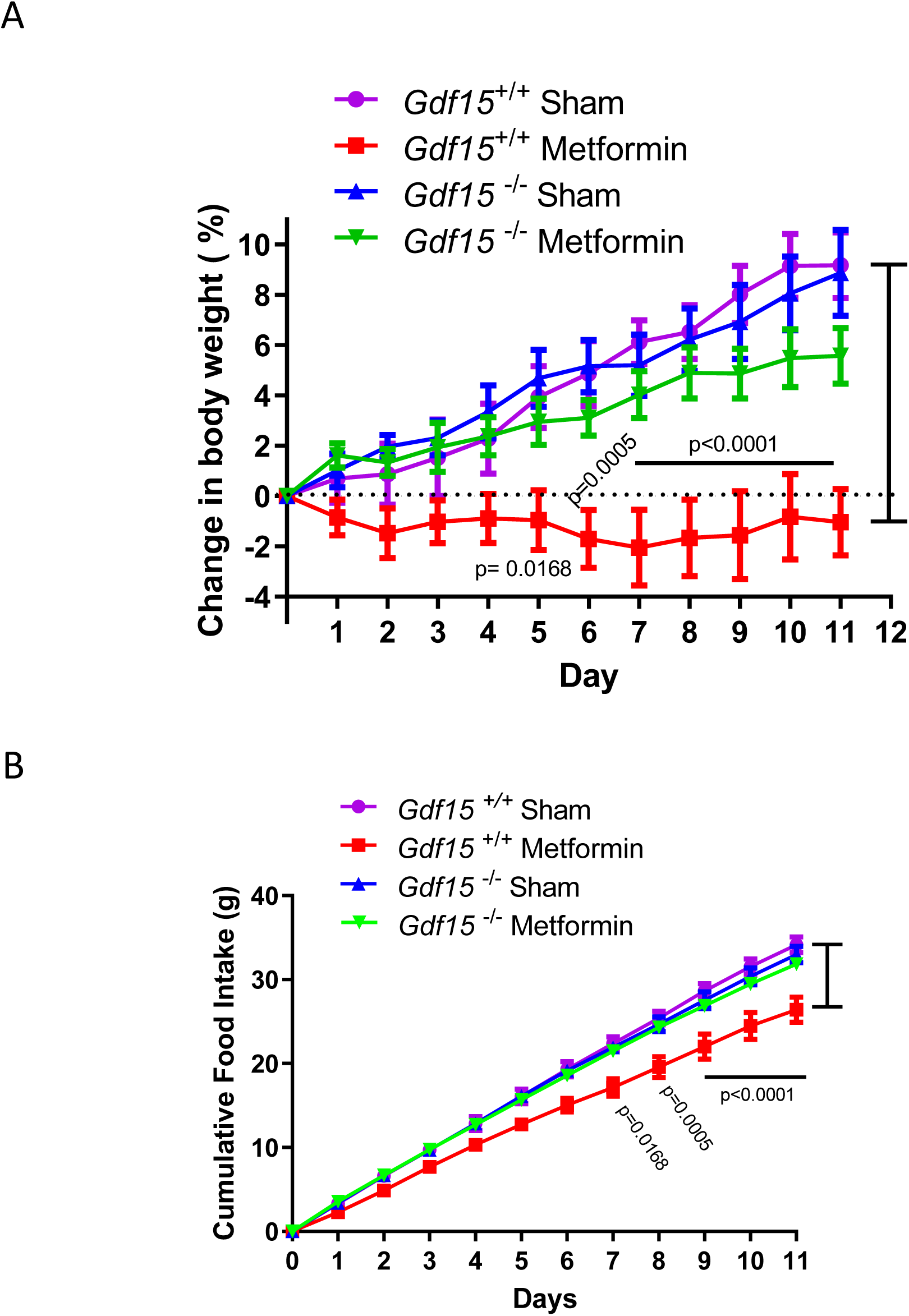
Response of second, independent cohort of high-fat diet fed *Gdf15* ^*+/+*^ and *Gdf15* ^*-/-*^ mice to metformin. A) Percentage change in body weight and B) cumulative food intake in *Gdf15 +/+* and *Gdf15* ^*-*^*/*^*-*^ mice on a high-fat diet treated with metformin (300mg/kg/day) for 11 days. Data are mean ± SEM (n=6/group, except *Gdf15* ^-/-^ sham= 8); *P* by 2-way ANOVA with Tukey’s correction for multiple comparisons.

**Supplementary Figure 4.**
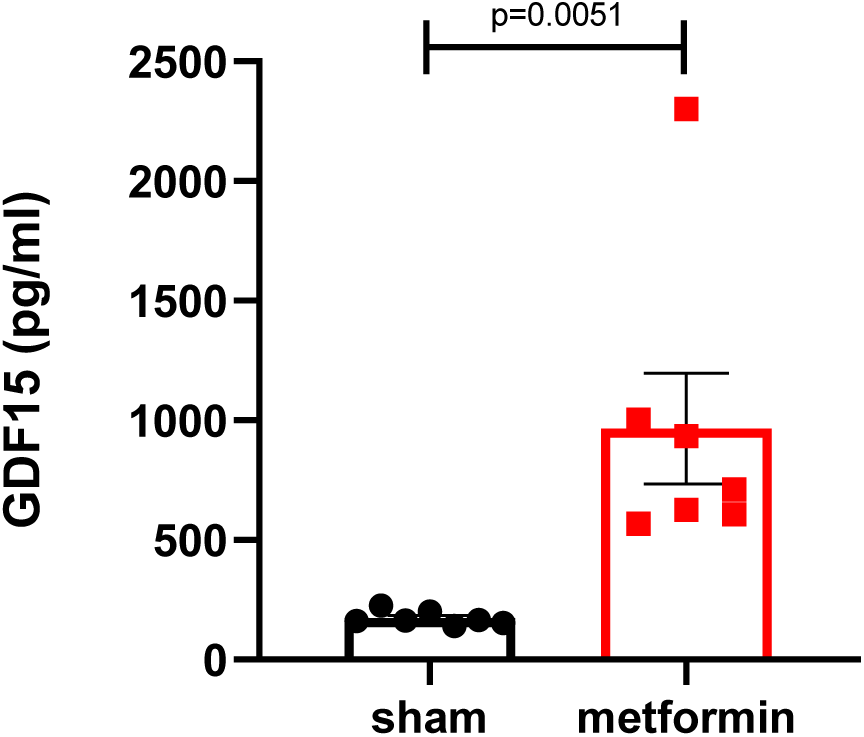
Circulating GDF15 levels in high-fat diet fed *Gdf15 +/+* mice after single oral dose of metformin (600mg/kg). Samples were collected 6 hours after dosing, data are mean ± SEM, (n=7/group), *P* by two-tailed *t*-test.

**Supplementary Figure 5.**
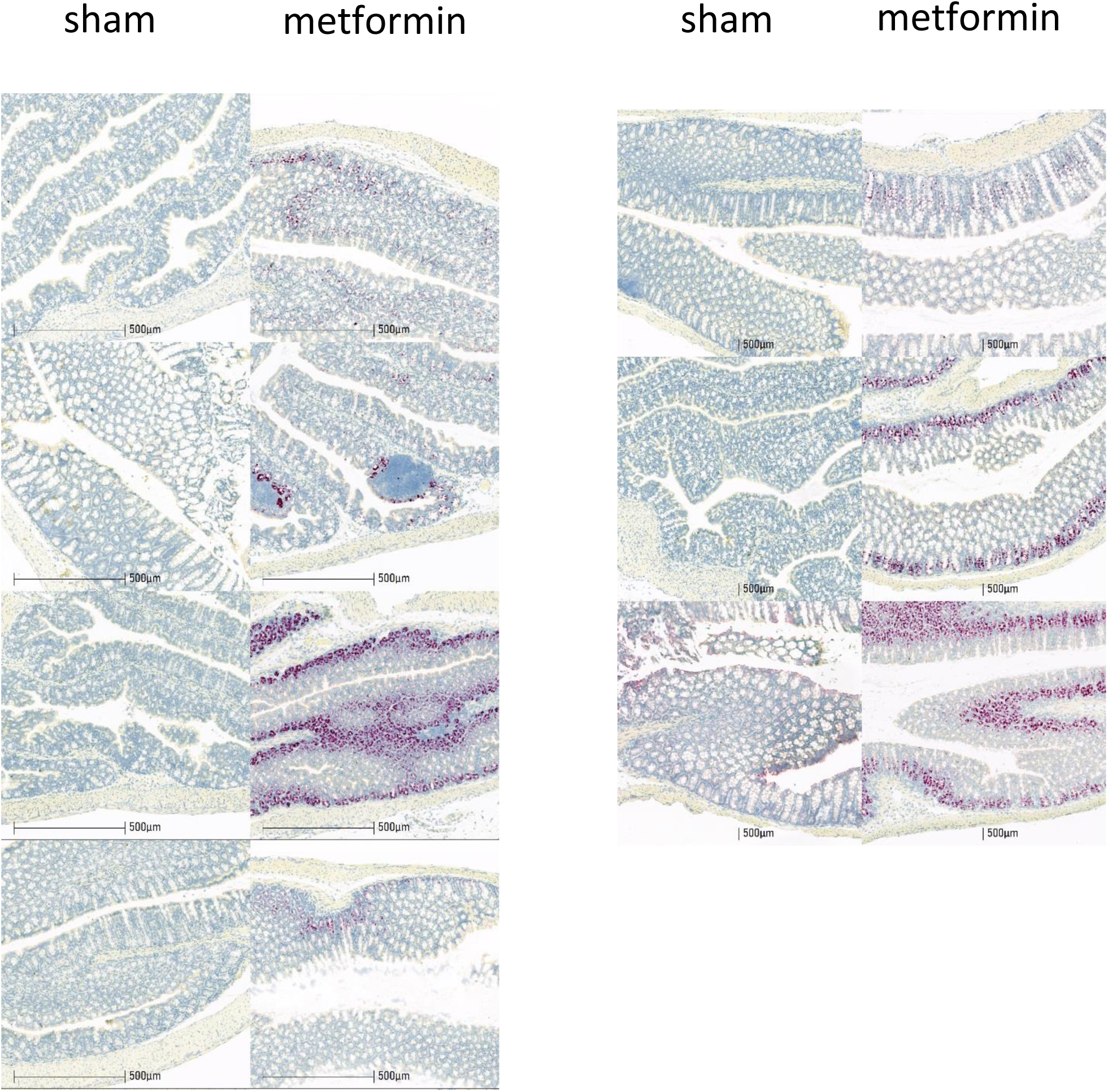
In situ hybridization for *Gdf15* mRNA expression (red spots) in all colons examined collected from high-fat fed wild type mice, 6 hrs after single dose of oral metformin (600mg/kg)(right side) or sham gavage (left side), n=7/group, mice as **Figure 4**.

**Supplementary Figure 6.**
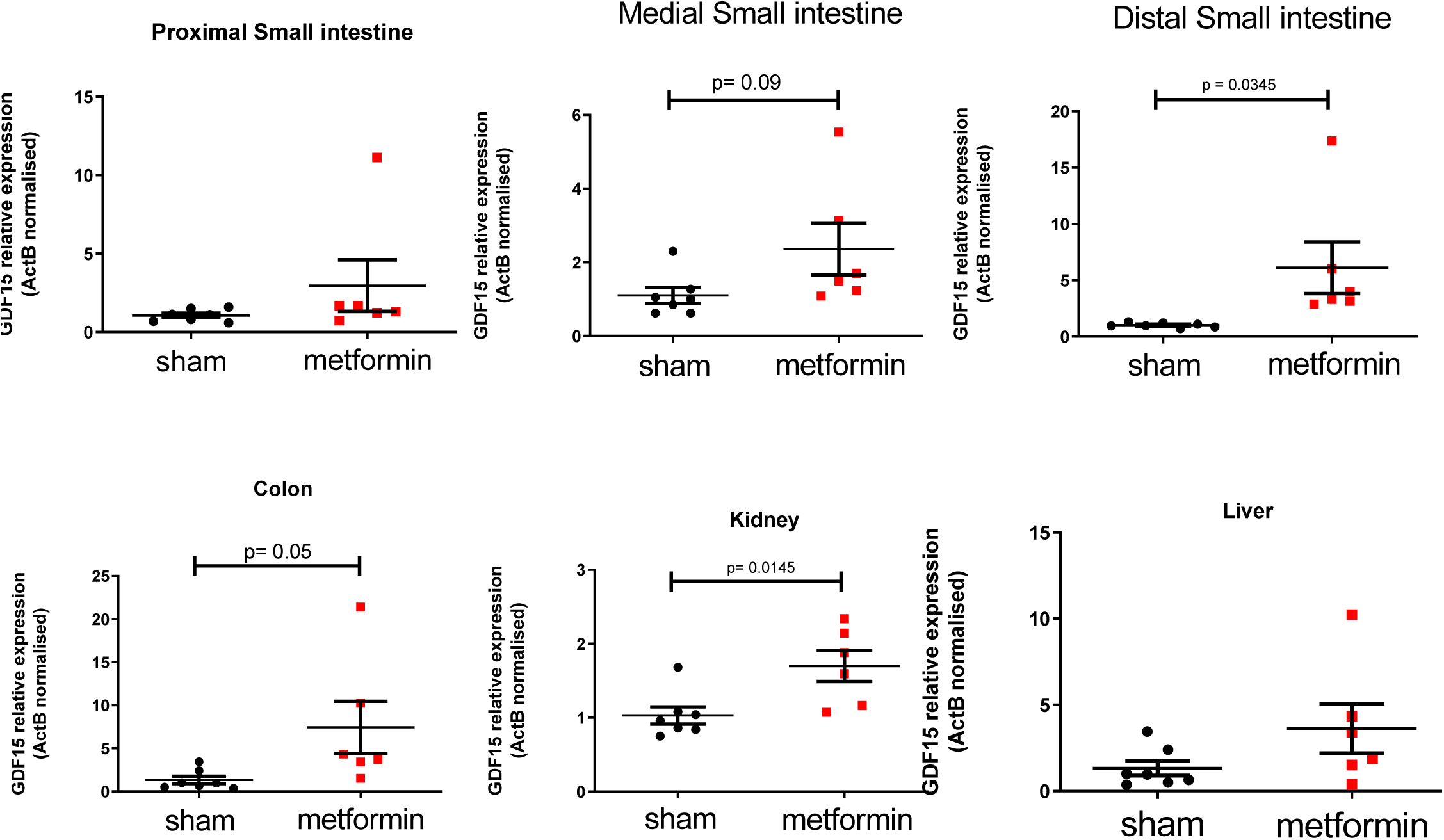
Analysis of *Gdf15* mRNA expression in tissue from high fat diet fed *Gdf15 +/+* mice given oral dose of metformin (300mg/kg) once daily for 11 days, normalised to expression levels of ActB (**see Figure 2A**). Data are mean ± SEM, n=6 metformin, n=7 sham, *P* by two tailed *t*-test.

**Supplementary Figure 7.**
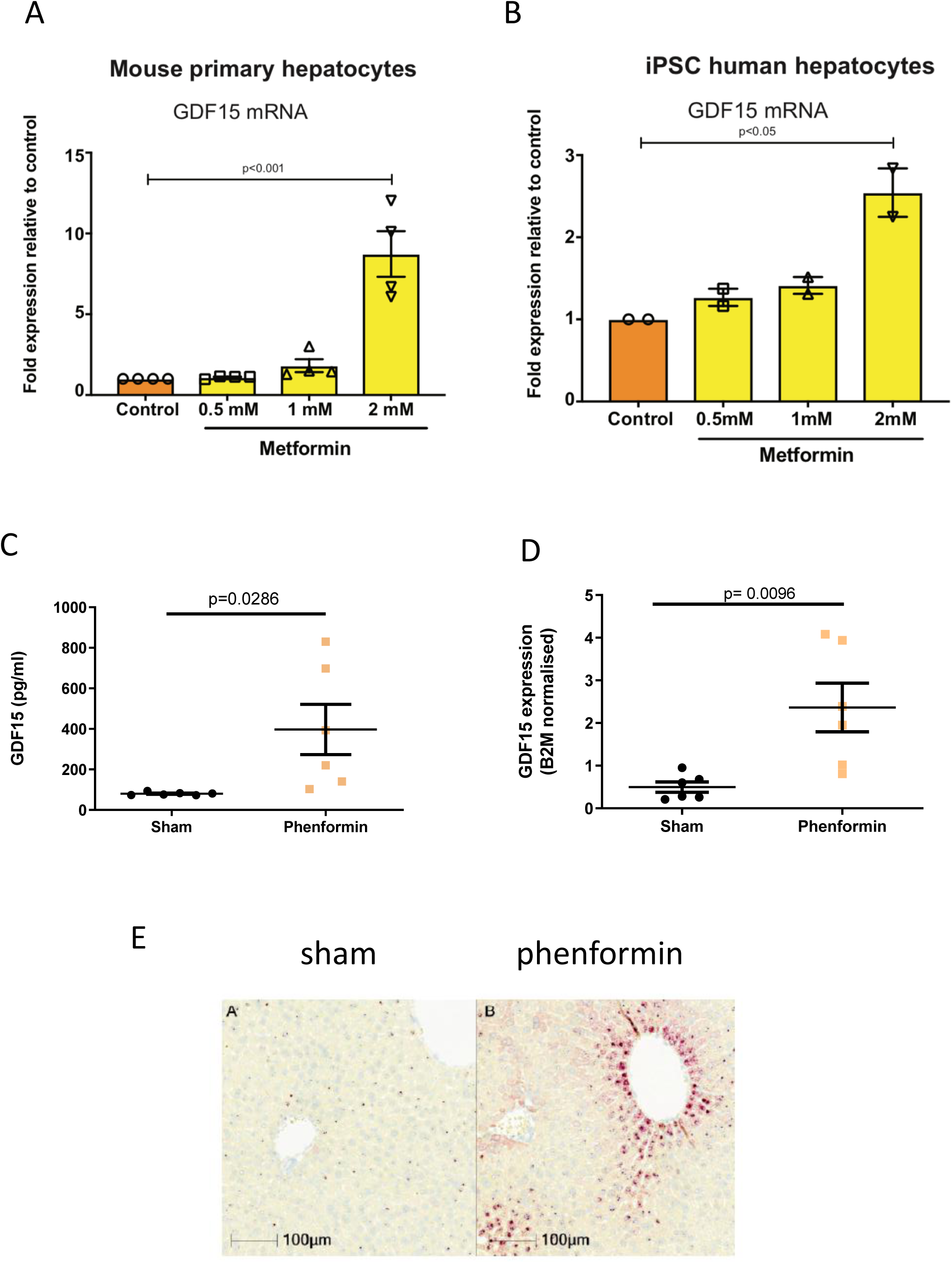
Hepatic GDF15 response to biguanides. GDF15 mRNA expression in (**A**) primary mouse hepatocytes or (**B**) human iPSC derived hepatocytes treated with vehicle control (Con) or metformin for 6 h. mRNA expression is presented as fold expression relative to control treatment (set at 1), normalised to HPRT and GAPDH gene in mouse and human cells, respectively. Data is expressed as mean ± SEM from four (A) and two (B) independent experiments. *P* by two-tailed *t*-test. (**C**) Circulating levels of GDF15 and (**D**) hepatic *Gdf15* mRNA expression (normalised to 2 microglobulin) in chow fed, wild type mice 4 hrs after single oral dose of phenformin (300mg/kg). Data are mean ± SEM, *P* by two tailed t-test. (**E**) Representative image of RNA scope analysis of fixed liver tissue derived from animals treated as described in (C) and (D).

